# Major Histocompatibility Complex (MHC) I-induced Endoplasmic Reticulum (ER) Stress mediates the secretion of pro-inflammatory muscle-derived cytokines

**DOI:** 10.1101/2022.06.29.496997

**Authors:** Anastasia Thoma, Kate E. Earl, Robert G. Cooper, Katarzyna Goljanek-Whysall, Adam P. Lightfoot

## Abstract

Major histocompatibility complex (MHC) I is an important component of intracellular antigen presentation. However, improper expression of MHC I upon the cell surface has been associated with several autoimmune diseases. Myositis is a rare acquired autoimmune diseases which targets skeletal muscle, and MHC I overexpression on the surface of muscle fibres and immune cell infiltration are clinical hallmarks. MHC I overexpression may have an important pathogenic role, mediated by the activation of the endoplasmic reticulum (ER) stress response. Given the evidence that muscle is a diverse source of cytokines, we aimed to investigate whether MHC I overexpression can modify the profile of muscle-derived cytokines and what role the ER stress pathway may play. Using C2C12 myoblasts we overexpressed MHC I with a H-2k^b^ vector in the presence or absence of salubrinal an ER stress pathway modifying compound. MHC I overexpression induced ER stress pathway activation and elevated cytokine gene expression. MHC I overexpression caused significant release of cytokines and chemokines, which was attenuated in the presence of salubrinal. Conditioned media from MHC I overexpressing cells induced *in vitro* T-cell chemotaxis, atrophy of healthy myotubes and modified mitochondrial function, features which were attenuated in the presence of salubrinal. Collectively, these data suggest that MHC I overexpression can induce pro-inflammatory cytokine/chemokine release from C2C12 myoblasts, a process which appears to be mediated in-part by the ER stress pathway.

## INTRODUCTION

The major histocompatibility complex (MHC) I is an integral component of antigen presentation processes and the immune response. Localised to the endoplasmic reticulum, MHC I plays a key role in the presentation of endogenous peptides upon the cell surface. Dysfunction in MHC I is a typical hallmark of several autoimmune diseases, with improper expression and overexpression of MHC I upon several cell and tissue types. One such disorder is myositis, a rare autoimmune disease which primarily affects skeletal muscle and is associated with overexpression of MHC I on the surface of muscle fibres (1). Patients with myositis typically present with symmetrical proximal muscle weakness and infiltration of CD4^+^ and CD8^+^ T cells into perifascicular and endomysial regions around non-necrotic fibres (1). The mechanisms underlying myositis remain poorly understood, though immune cell mediated and immune cell independent processes (e.g., ER stress) likely both play a role (2-4). Overexpression of MHC I on and within muscle cells in myositis patients are associated with activation of the ER stress pathway within muscle cells, in both murine myositis models and in human disease (5). Markers of ER stress pathway activation (e.g., Grp75 and Grp94) co-localise within muscle fibres staining positive for MHC I in myositis patient biopsies (6). Moreover, activation of the ER stress response is known to activate inflammatory pathways, such as nuclear factor kappa B (NF-κB), a pleiotropic transcription factor capable of activating many downstream pathways to initiate and propagate inflammation (7).

Skeletal muscle is increasingly recognised as an endocrine organ capable of secreting a diverse range of peptides and proteins, components of the secretome (8). The issue of muscle-derived cytokine production and release has mostly been examined in the context of exercise (9). Studies of diagnostic muscle biopsies from myositis patients have demonstrated elevated gene expression of several cytokines and chemokines including: CCL3, CCL4, CCL5, IL-6, IL-15 and CCL2 (10, 11). Moreover, there has been increased interest in better understanding the potential (patho-)physiological roles of muscle-derived cytokines (myokines) in the inflammatory myopathies (12). However, the source of biopsy-localised cytokines remains poorly understood, where muscle and infiltrating immune cells will both likely harbour these factors.

Given the established involvement of MHC I upregulation in myositis and association with ER stress, the ER overload response (EOR) and nuclear factor kappa B (NF-kB) activation (5) we have used a cell model to determine whether MHC I overexpression can induce cytokine production and release from C2C12 myotubes, and whether compounds which affect the ER stress response (salubrinal) can augment these changes. Furthermore, we tested whether factors secreted from MHC I overexpressing cells can elicit changes in normal untreated C2C12 cells.

## METHODS

### Cell culture, treatment, and transfection

C2C12 myoblasts (13) were grown in culture under standard conditions in Dulbecco’s modified eagles’ medium (DMEM) growth media supplemented with 10% foetal calf serum, 2mM L-glutamine, then differentiated into mature myotubes over 7 days in media supplemented with 2% horse serum (HS) (14). To upregulate MHC I levels, myotubes were transfected with 1µg of the MHC I (H2-k^b^) overexpression plasmid using Lipofectamine™ 2000 reagent for a period of 6 hours (Invitrogen, USA), both without and with the addition of the ER stress blocking agent, salubrinal (1µM), with a green fluorescent protein (GFP) plasmid used as an additional control, transfection efficiency was between 40-70% for each construct (15). Cells and conditioned media were harvested 18 hours later in and stored at -80°C pending subsequent analyses. The plasmid used (MDH1-PGK-H2-Kb^2^) was a gift from Chang-Zheng Chen (Addgene plasmid #17852) (16). Pharmacological induction of ER stress in C2C12 myotubes was achieved by treating cells with Tunicamycin (0.1µg/ml) for 24 hours (17).

### Conditioned media experiments and microscopy

Normal C2C12 myotubes were grown in culture as previously described, and following which, were treated for five days with conditioned media harvested from C2C12 myotubes, transfected with H2-k^b^, GFP with or without the presence of salubrinal. After five days of treatment with conditioned media, cells were imaged to assess myotube morphology/diameter. Images were acquired using a Nikon Eclipse TE 2000 microscope (Nikon Instruments B.V, UK), and analysed using ImageJ software. Following imaging, cells were harvested in ice-cold PBS and stored at - 80°C for gene expression analyses by qPCR.

### RNA extraction, cDNA synthesis and qPCR

RNA was extracted from myotubes using the TRIzol method, purified using Genejet RNA kit (Thermo Fisher Scientific, UK), and cDNA synthesised using the iScript first strand kit (Bio-Rad, Heracles, USA). Quantitative PCR was performed with SYBR green (Roche Diagnostics, UK) and analysed by the 2^-^ΔΔct method (18). The primer sequences used are detailed in (**Table 1**). The choice of the analysed ER stress pathway genes (i.e., PERK, IRE1 and ATF6) was made because these represent the three main branches of the ER stress pathway thus permitting analysis of the whole pathway (19). In addition to examining Grp78, which has been previously examined in myositis, we also examined XBP-1, which is recognised as having a cytokine-regulating role (5, 20).

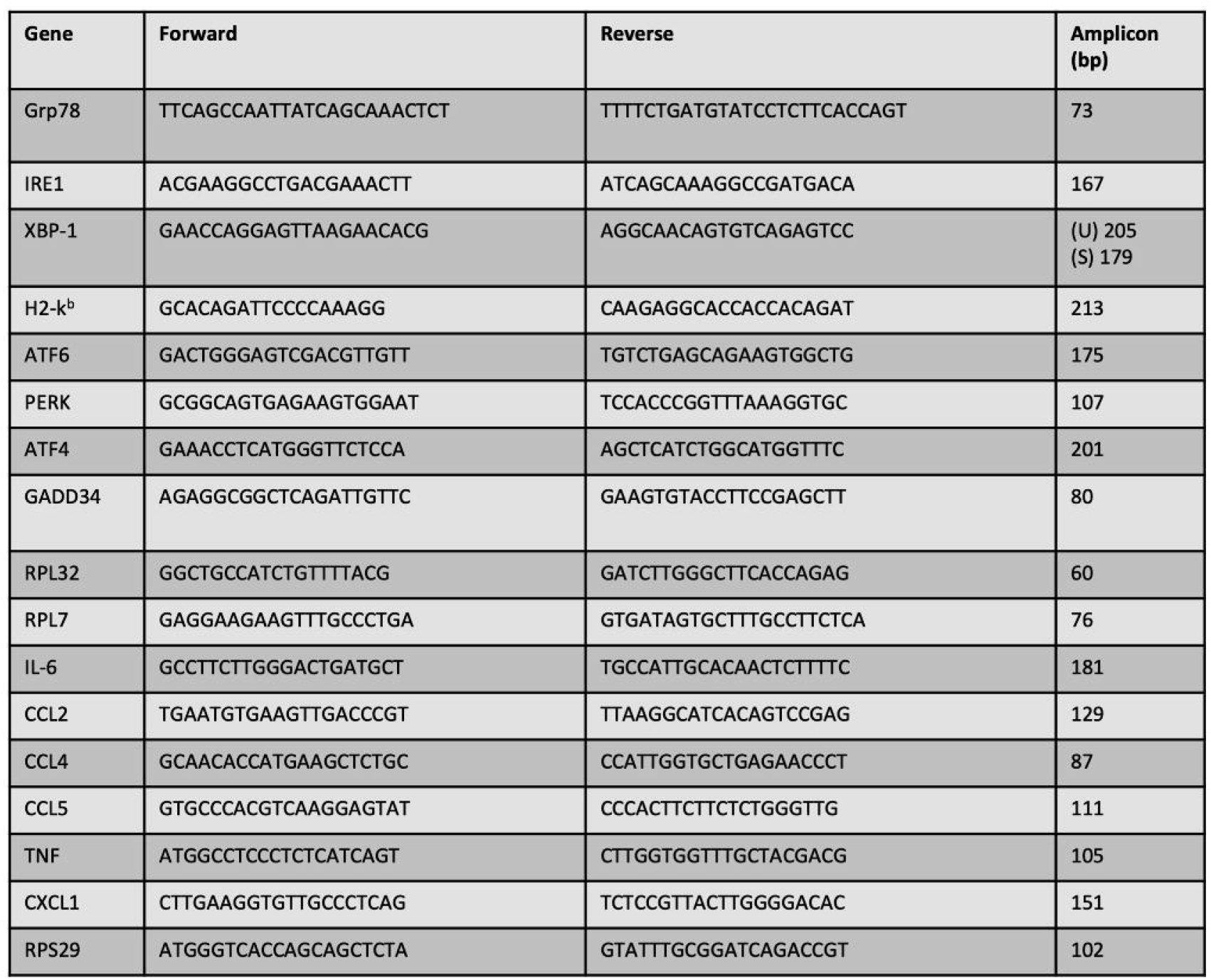

### Multiplex cytokine analysis

Conditioned media from transfected cells were analysed for cytokine content by multiplex bead analysis with antibodies specific to IL-6, CXCL1, TNF-α, CCL2, CCL4 and CCL5 using a Bioplex-200 platform in accordance with the manufacturers protocol (Bio-Rad, Heracles, USA). This choice of cytokine and chemokine targets was not designed to be comprehensive but based instead on the results from our previous work, which identified several specific cytokines/chemokines released from C2C12 myotubes (14).

### Chemotaxis Assay

The murine CD4^+^ T lymphocyte cell line (BW5147.3) used in the chemotaxis assay studies was obtained from the American Tissue Culture Collection (ATCC), and cultured in suspension, in growth media comprising DMEM containing 10% HS, 2mM L-glutamine. The ability of conditioned media from H2-K^b^ transfected cells (± salubrinal) to chemoattract murine CD4^+^ T lymphocytes was assessed using a CytoSelect 96-well Cell Migration Assay, with a lymphocyte specific 5µM pore filter (Cell Biolabs Inc), in accordance with the manufacturers protocol.

### Mitochondrial function analyses

C2C12 cells were cultured and exposed to conditioned media as previously described, in 8-well seahorse XFp plates. Real time oxygen consumption in cells was measured using a Seahorse XFp extracellular flux analyser, using the XFp mito stress test (Agilent Technologies, Manchester, UK), performed in accordance with manufacturer’s instructions (17).

### Statistical analysis

Data are presented as mean ± SEM; statistical analyses of data were undertaken using analysis of variance (ANOVA) with Tukey *post hoc* test and Kruskal-Wallis test, following shapiro-wilk assessment of normality, using GraphPad Prism 8.

## RESULTS

### Overexpression of MHC I resulted in activation of the ER stress pathway

Transfection of myotubes with the MHC I (H2-k^b^) vector in the presence or absence of salubrinal resulted in significant increase in MHC I gene expression (**Figure 1A**). MHC I overexpression resulted in activation of several ER stress pathway components examined, i.e., *IRE1, PERK* and *ATF6*, as well as the chaperone *Grp78*, irrespective of the presence of salubrinal (**Figure 1B - F**).

**Figure 1:**
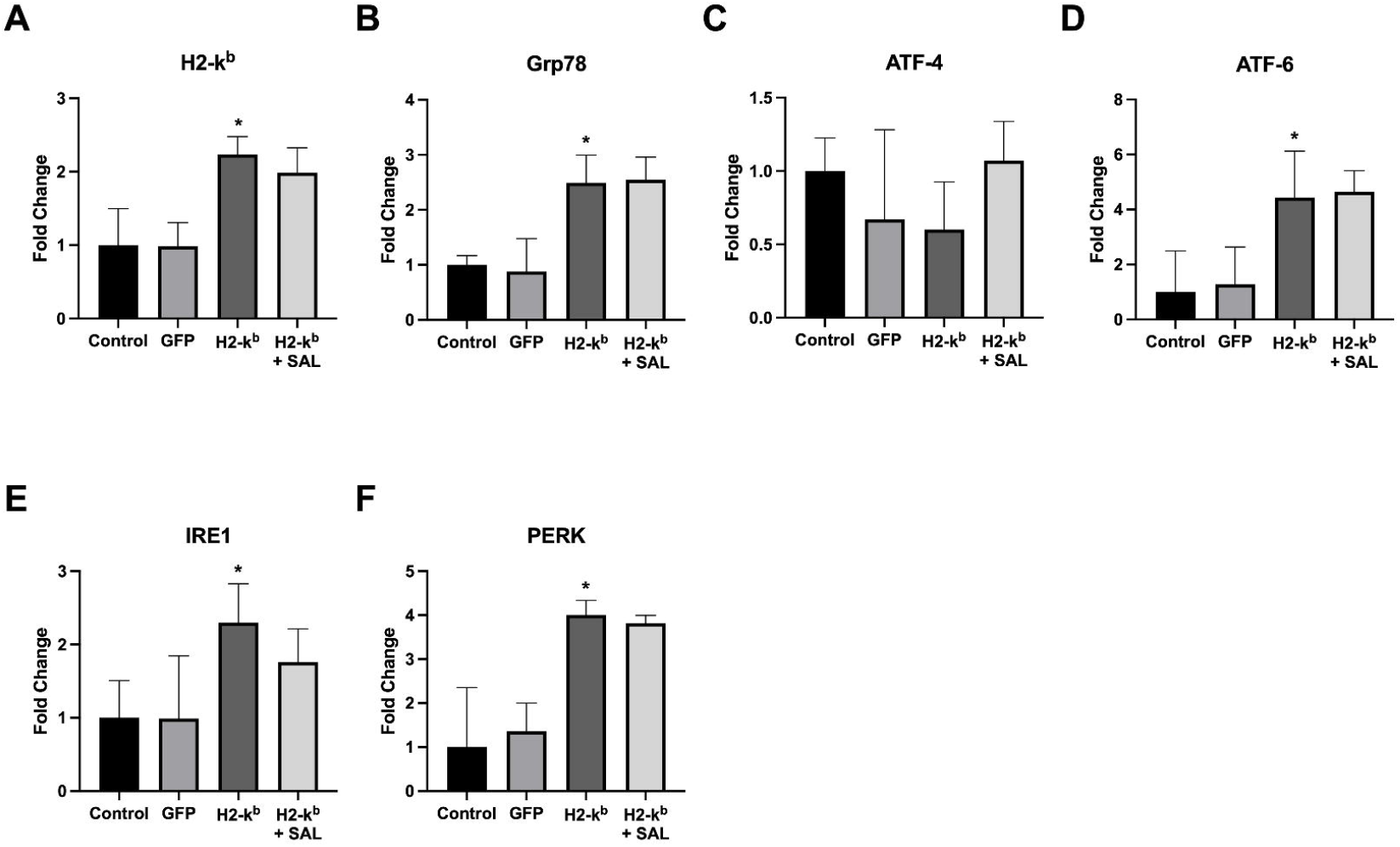
H2-k^b^ (MHC I) **(A)**, Grp78, ATF-4, ATF-6, IRE1, PERK **(B)**, gene expression levels in C2C12 myotubes transfected with the GFP vector, H2-k^b^ vector with or without the presence of salubrinal. Data are presented as mean ± SEM (n=3-6), p≤ 0.05*.

### MHC I overexpression resulted in increased cytokine gene expression and ER stress-induced release of chemotactic myokines

Overexpression of MHC I in C2C12 myotubes resulted in significant increases in cytokine gene expression (**Figure 2A**) and cytokine release into the culture media, including of: IL-6, CXCL-1, CCL2, CCL4 and CCL5. Salubrinal treatment resulted in a reduced release of IL-6, CCL2, CCL4 and CCL5, but not of CXCL-1 (**Figure 2B**). In contrast, TNF-α mRNA gene expression was undetectable, and cytokine release not increased in response to MHC I overexpression (**Figure 2B**). Conditioned media from MHC I overexpressing cells induced significant chemotaxis of CD4^+^ T lymphocytes, an effect significantly reduced by salubrinal (**Figure 3**). Treatment of C2C12 myotubes with tunicamycin induced the upregulation in the gene expression (**Figure 4A**) and protein release of IL-6, CXCL1, CCL2 and CCL5 from normal C2C12 cells (**Figure 4B**).

**Figure 2:**
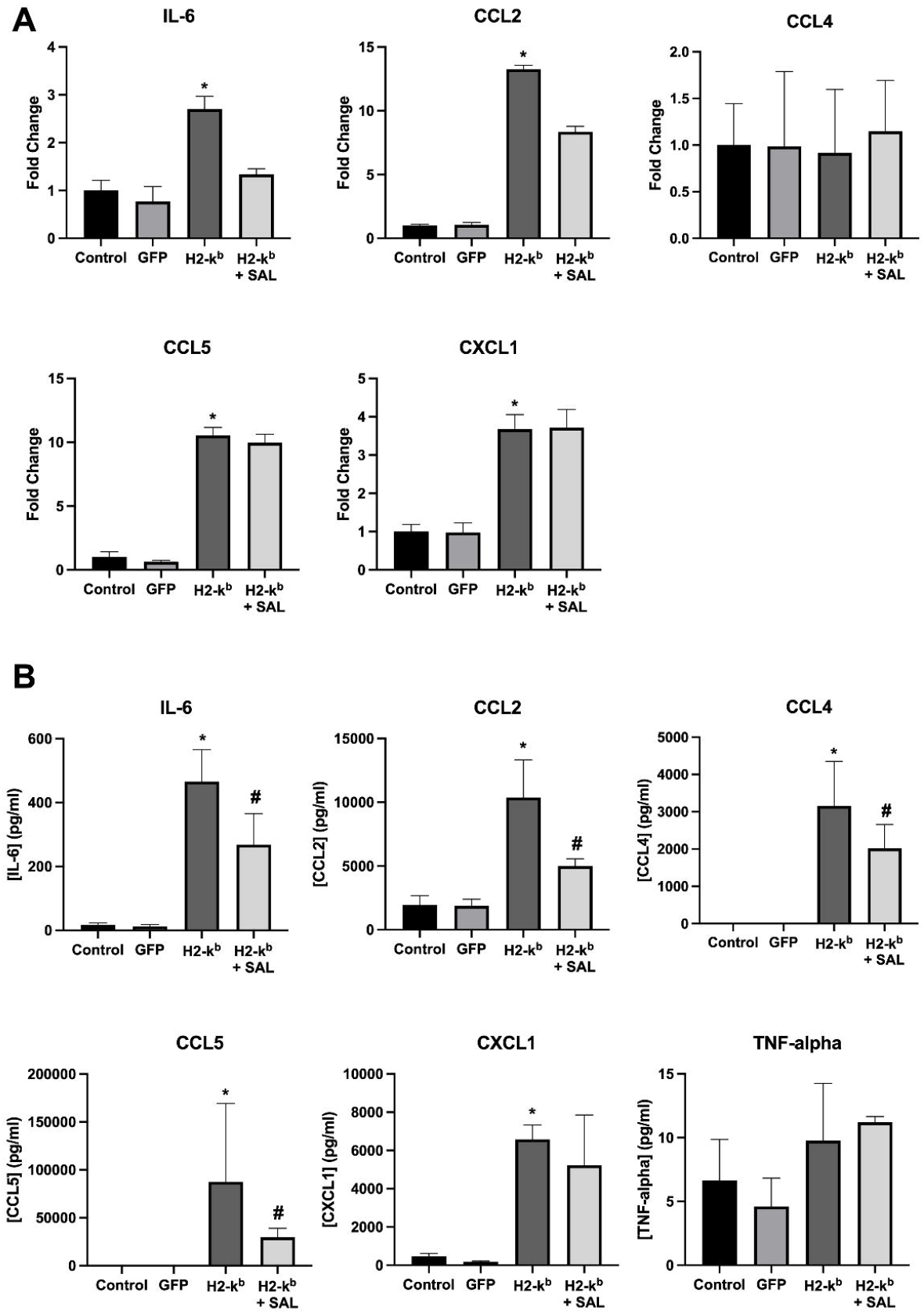
IL-6, CCL2, CCL4, CCL5, CXCL1 **(A)**, gene expression levels and protein levels (**B**) released from C2C12 myotubes transfected with the GFP vector, H2-k^b^ vector with or without the presence of salubrinal. Data are presented as mean ± SEM (n=3-6), p_≤_ 0.05*.

**Figure 3:**
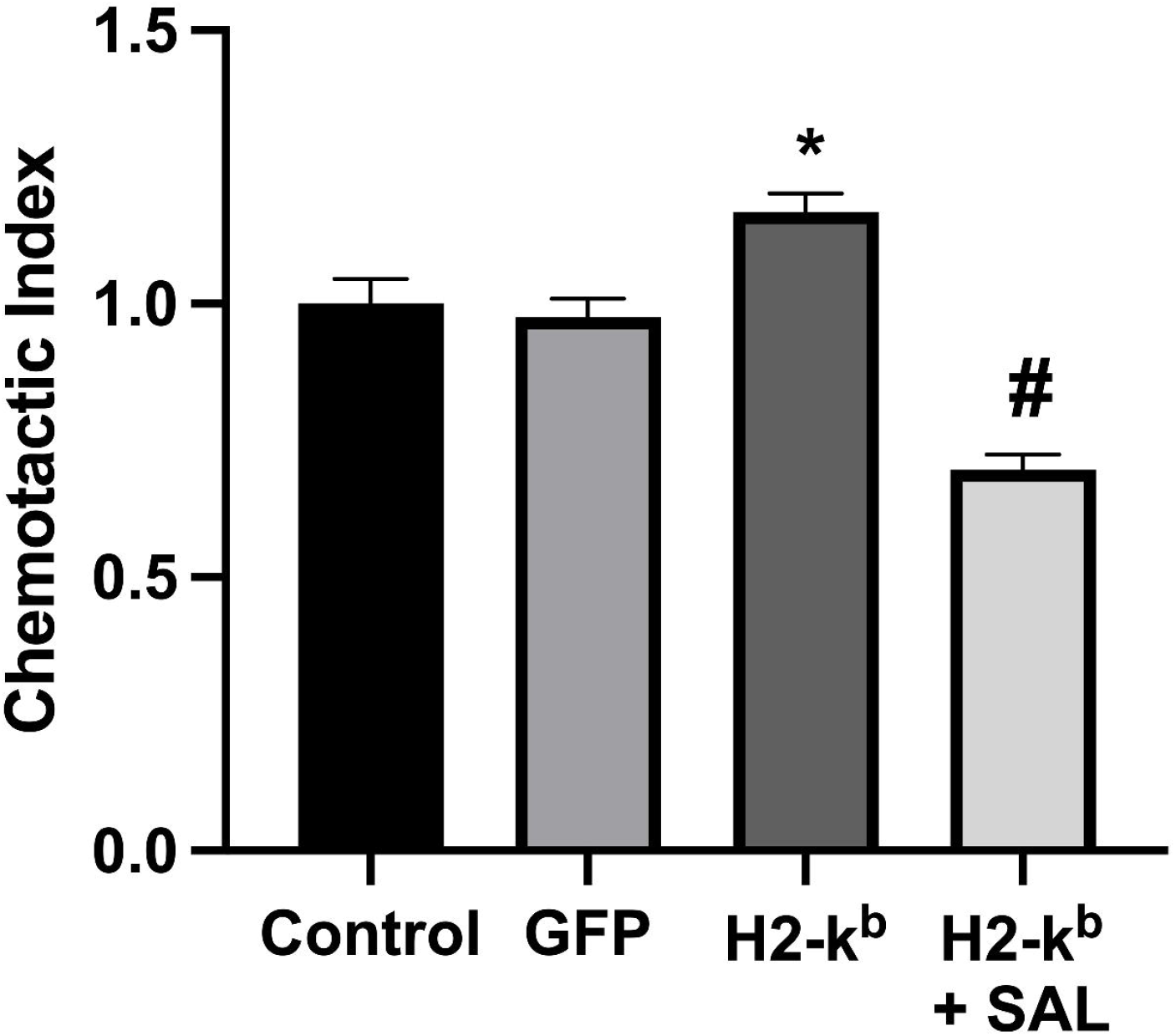
T-cell chemotaxis in response to cell culture media taken from C2C12 myotubes transfected with the H2-kb vector with or without the presence of salubrinal. Data are presented as mean ± SEM (n=3-6), p≤ 0.05*.

**Figure 4:**
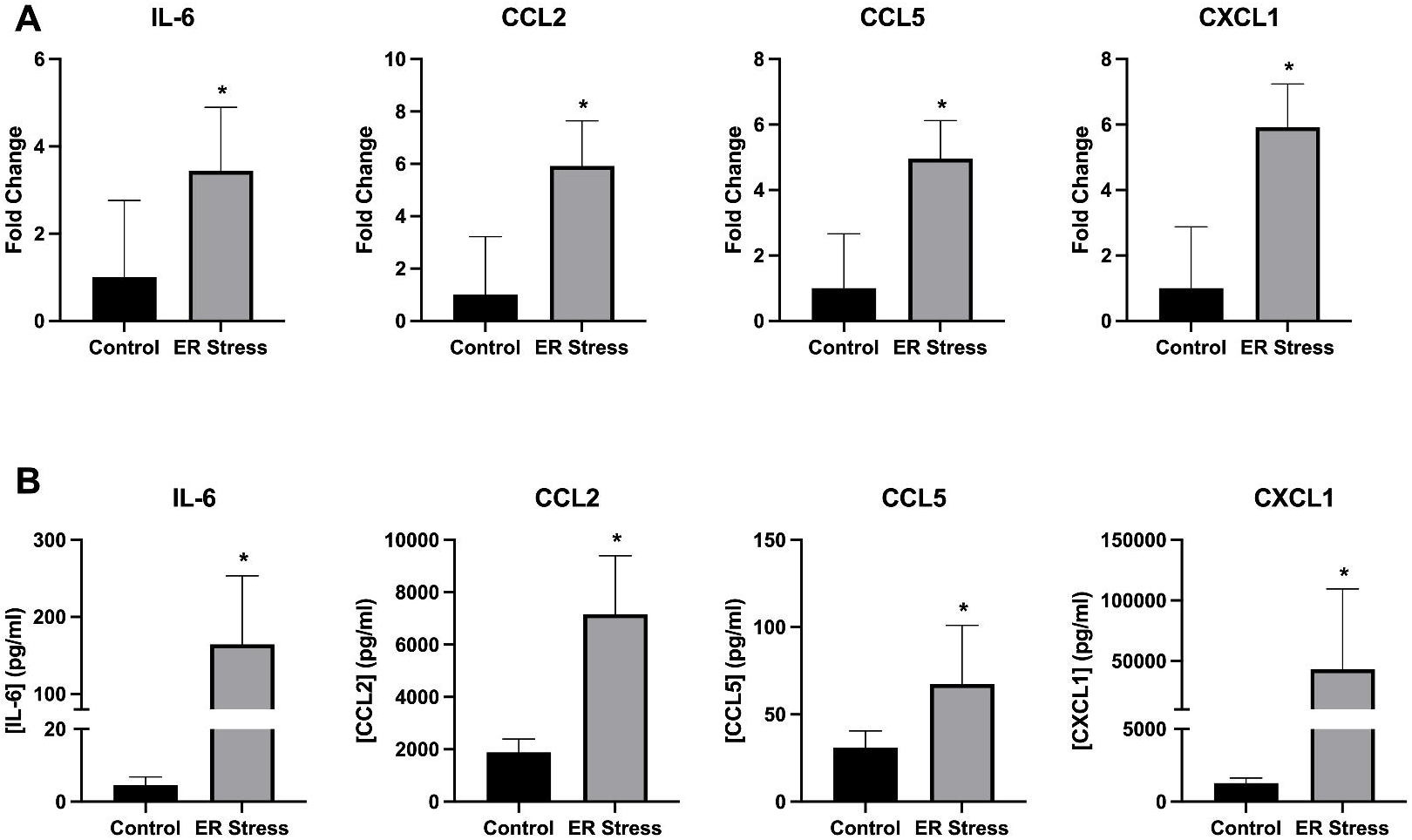
Gene expression **(A)** and released protein levels **(B)** of IL-6, CXCL1, CCL2 and CCL5 from C2C12 myotubes treated with 0.1µg/ml of the ER stress-inducing compound tunicamycin for 24 hours. Data are presented as mean ± SEM (n=3-6), p≤ 0.05*.

### Conditioned media from MHC I overexpressing cells induced atrophy and mitochondrial dysfunction in untreated C2C12 cells

C2C12 cells incubated with conditioned media from MHC I overexpressing cells demonstrated a significant reduction in diameters (**Figure 5-E**). The latter was clearly less marked in myotubes, which were incubated in conditioned media from cells overexpressing MHC I in the presence of salubrinal (**Figure 5A-D**). Conditioned media from MHC I overexpressing cells induced a significant up-regulation in the gene expression of two key genes which have been associated with muscle atrophy (21), *Atrogin-1* and *MuRF-1* (**Figure 5F-G**). *MuRF-1* expression was reduced in cells treated with conditioned media from MHC I transfected cells in the presence of salubrinal. Conditioned media from MHC I overexpressing cells induced a decline in oxygen consumption (**Figure 6A**), specifically increased proton leak (**Figure 6D**) from mitochondria and significant reduction in non-mitochondrial respiration (**Figure 6E**).

**Figure 5:**
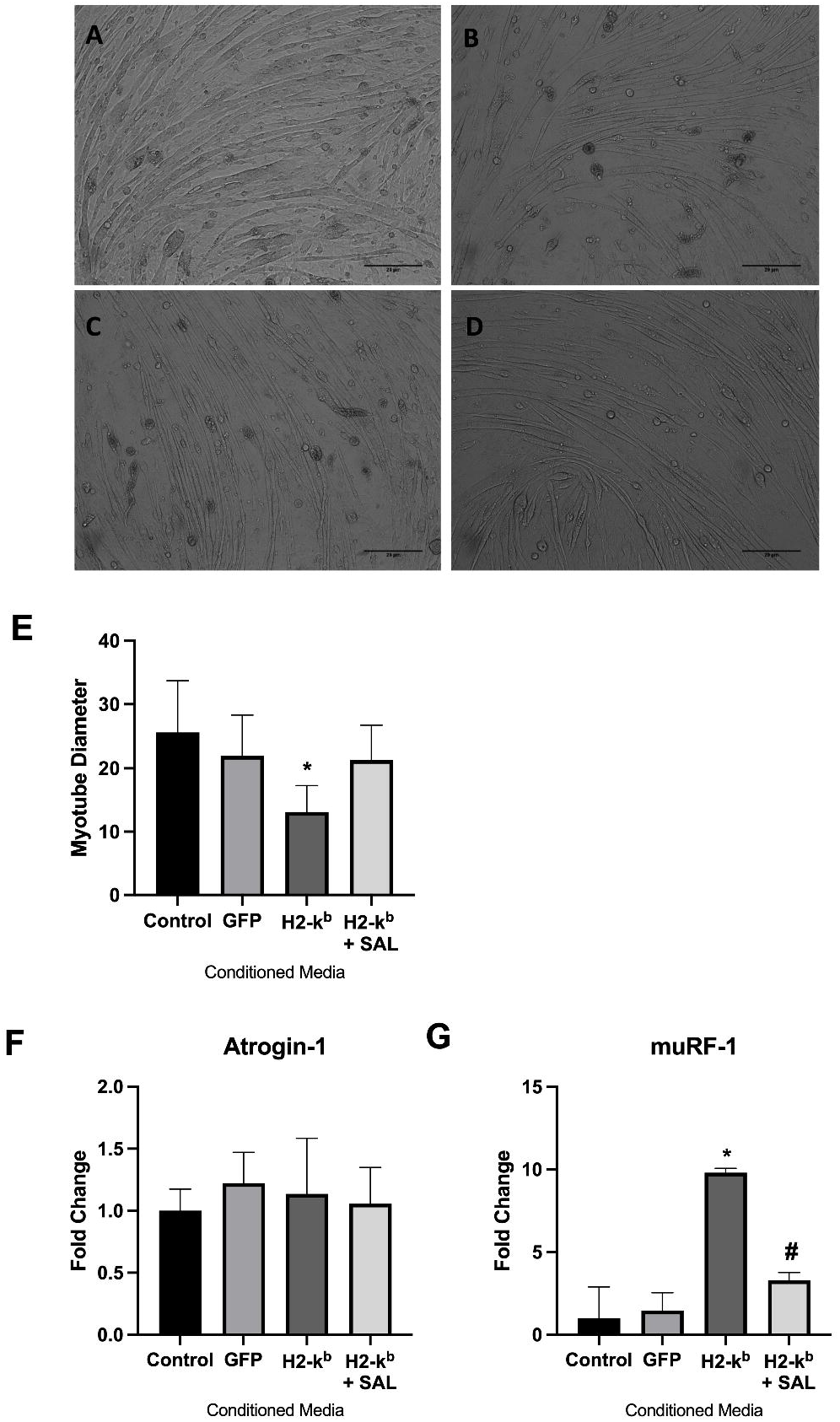
Representative images (x10 magnification) of normal C2C12 myotubes treated with conditioned media taken from control **(A)**, GFP transfected **(B)**, H2-k^b^ transfected **(C)** and H2kb transfected with salubrinal (1µM) **(D)** C2C12 myotubes. Quantified myotube diameter **(E)** and gene expression levels of *Atrogin-1* **(F)** and *MuRF-1* **(G)** in normal C2C12 myotubes treated with conditioned media taken from C2C12 myotubes transfected with control, GFP, H2-k^b^ or H2-k^b^ with salubrinal. Data are presented as mean ± (n=4) p_≤_0.05*.

**Figure 6:**
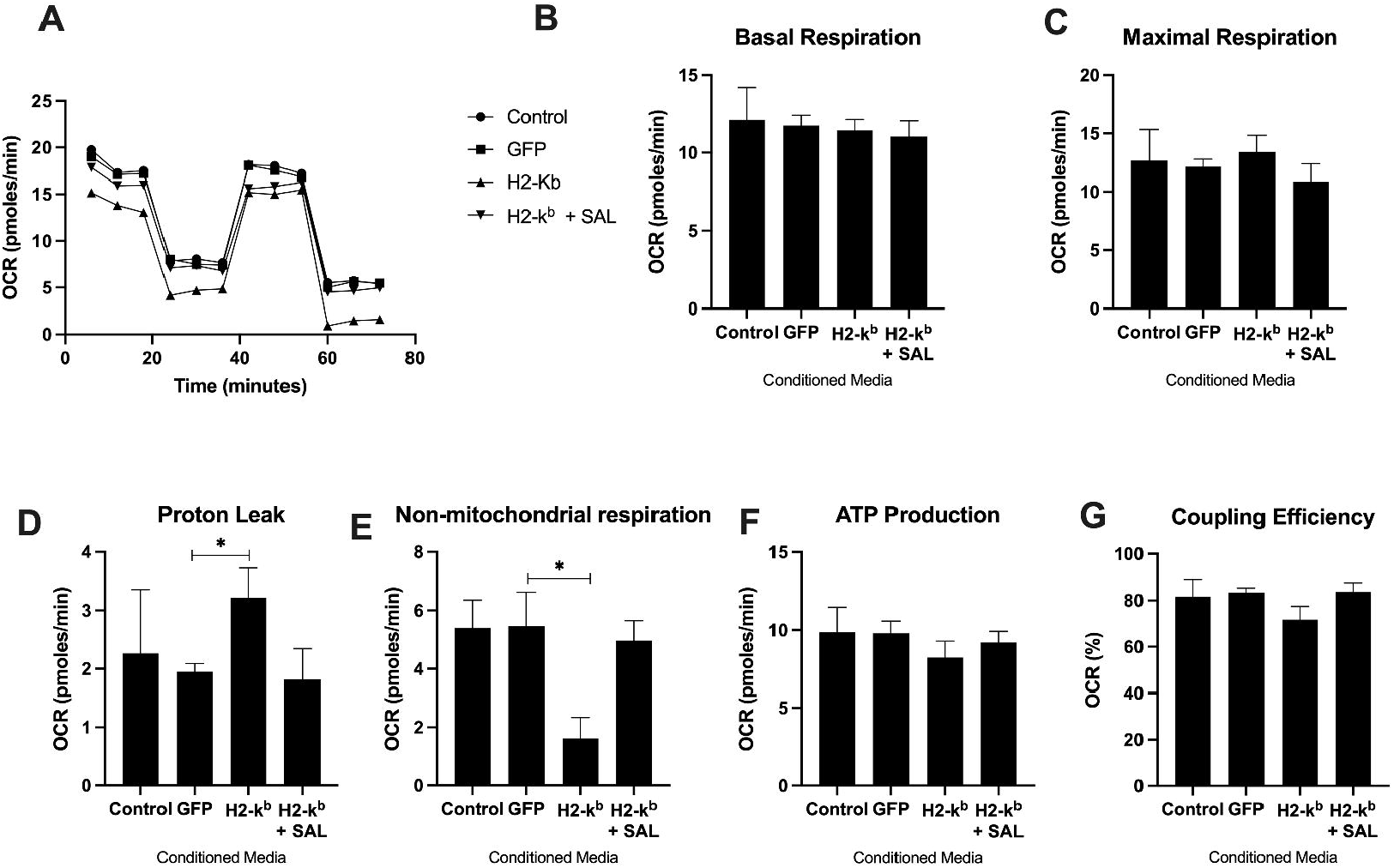
Mitochondrial function measures, showing overall oxygen consumption rate **(A)**, basal respiration **(B)**, maximal respiration **(C)**, proton leak **(D)**, non-mitochondrial respiration **(E)**, ATP-linked respiration **(F)** and coupling efficiency **(G)** in normal C2C12 myotubes treated with conditioned media taken from C2C12 myotubes transfected with control, GFP, H2-k^b^ or H2-k^b^ with salubrinal. Data are presented as mean ± (n=4) p_≤_0.05*.

## DISCUSSION

The aim of this study was to investigate the impact of overexpressing MHC I in skeletal muscle cells on cytokine secretion and the potential role of ER stress modifying compound salubrinal. Further, to assess the impact of secreted factors on normal C2C12 cells.

Our findings provide insight into the impact of MHC I overexpression in skeletal muscle cells, suggesting that MHC I-induced ER stress may play a role in cytokine expression and release from C2C12 myotubes. The observation of increased Grp78 expression in MHC I overexpressing cells validates previous evidence suggesting that ER stress activation is downstream of MHC I upregulation and is somewhat analogous to observations seen in certain *in vivo* murine myositis models as well as human myositis (5). The demonstrated increased expression of key ER stress pathway receptors (i.e., IRE1 and ATF6) following upregulation of MHC I is novel, and perhaps suggests broad activation of the pathway. Moreover, the pharmacological induction of ER stress, showed similar patterns of muscle-derived inflammatory cytokine expression and release to that of the MHC I overexpression.

Many cytokines have been detected in muscle tissue from myositis patients and the current data suggest that muscle may also be a source of some of the chemokines and cytokines detectable in muscle biopsies from myositis patients (22-24). The cytokines released by myotubes in our study do not reflect the full spectrum of cytokines detected in myositis patient muscle biopsies. Equally, infiltrating immune or vascular endothelial cells may also be significant sources for these and other cytokines detected in myositis patient biopsies (22, 25). TNF-α was not induced by MHC I overexpression in our cell model, given that this cytokine has so frequently and specifically been implicated in contributing directly to muscle dysfunction in a considerably variety of systemic diseases (7). TNF-α gene expression has, for instance, been detected in the sarcopenic skeletal muscle of older individuals, and co-localised to type I muscle fibres and in myositis patient biopsies (26-28). However, there is little evidence to suggest that skeletal muscle secretes TNF-α, though transcript levels are detectable in some cellular models, albeit at very low levels (29). The expression of both CXCL and CCL chemokines in our model is consistent with similar *in vitro* studies, following TNF-α stimulation (14). CCL2 (MCP-1) has been reported to be associated with T-cells in non-necrotic muscle fibres of patients with PM and sIBM (30). Moreover, the expression of CCL4 and CCL5, along with their requisite receptors has been reported in biopsies from patients with inflammatory myopathies (31). The function of these potential myokines remains poorly understood.

The release of IL-6 downstream of MHC I is an interesting observation, there is a significant juxtaposition towards the function of IL-6 in muscle. Studies have reported IL-6 can induce muscle atrophy (32), however, IL-6 has been shown to be protective in a murine model of myositis (33). Despite these somewhat paradoxical observations, the pursuit of anti-IL-6 therapies in inflammatory myopathies (and wider autoimmune diseases) is highly prevalent (34, 35).

The suppressed cytokine release in the presence of salubrinal is likely a consequence of its role in translational attenuation, through preventing the dephosphorylation of eiF2-alpha (15). Studies in Parkinson’s disease have reported similar attenuation of the circulating levels of cytokines IL-1β, IL-6 and TNF-α in LPS treated mice (36). Our findings support the hypothesis that, in myositis, muscle-derived cytokines may play an important local influential role to effect neighbouring fibres, and in creating a milieu, which could contribute to disease pathogenesis (37).

Given that myofibre atrophy and ER stress are characteristic hallmarks of myositis (38) our findings suggest that muscle-derived factors could play a pathogenic role in these processes. For instance, myokines released following upregulation of MHC I, could exert pro-atrophic effects in neighbouring muscle fibres. In our model, myotube atrophy was lessened in cells treated with conditioned media derived from MHC I overexpressing cells when these were treated with salubrinal. Surprisingly, *atrogin-1* gene expression levels remained elevated, despite incubation with conditioned media derived from MHC I overexpressing cells treated with salubrinal. This may be due to the fact that *MuRF-1* and *atrogin-1* have different regulators.

Mitochondrial dysfunction has been described in both experimental and human models of myositis; moreover, mitochondrial dysfunction as been associated with ER stress (39, 40). The mechanisms responsible are overall poorly characterised, however, exogenous factors such as type I interferon have been heavily implicated (39). Muscle precursor cells have been reported as a potential source of type I interferon in myositis (41). Moreover, recent data has shown primary cells from myositis patients display an intrinsic deficit in mitochondrial function (42). Our data showed a reduction in overall OCR and evidence of mitochondrial dysfunction through increased proton leak (a marker of mitochondrial damage) in cells treated with conditioned media from MHC I overexpressing cells, where media from salubrinal treated cells ameliorated these changes. Furthermore, the decline in non-mitochondrial respiration suggests perhaps a deficiency in broader metabolic processes. Collectively, our data suggest that secreted factors from disease phenotype cells may affect function in neighbouring cells/fibres through augmenting the local milieu and compound the intrinsic mitochondrial deficits that have previously been reported (42).

The ER stress pathway is a cellular mechanism highly conserved across a wide range of eukaryotes (43). Thus, although our experiments were performed in a murine cell line only, we anticipate the mechanisms reported here would be conserved across species. Previous studies in a murine model of antibody-induced inflammatory arthritis reported that reduction of ER stress by salubrinal decreased the levels of induced joint inflammation (44).

There are clear limitations to this study, the applicability of these findings towards human disease remains unknown, the use of a murine cell line is indeed a limiting factor – future studies on human cells are needed. Equally, the overexpression of MHC I is an artificial induction in our model, in human myositis, there a likely an array of driving factors, further the magnitude of MHC I overexpression in our model may not be truly reflective of the phenotype seen in patients. Even though our findings provide the foundation for the role of MHC I-mediated myokine release, further experiments on patient-derived cells are needed to contextualise these findings.

## Funding

This study was funded by the University of Liverpool, Myositis UK and The Manchester Metropolitan University.

### ABBREVIATIONS

ATF: activating transcription factor
CK: creatine kinase
DM: dermatomyositis
eif2_α_: eukaryotic translation initiation factor 2 alpha
ER: endoplasmic reticulum
Grp: glucose-regulated protein
IRE1: inositol-requiring enzyme 1
MHC: major histocompatibility complex
NF_-κ_B: nuclear factor kappa B
PERK: protein kinase RNA-like endoplasmic reticulum kinase
sIBM: sporadic inclusion body myositis
XBP-1: x-box binding protein 1
MuRF-1: muscle RING-finger protein-1
PM: polymyositis.

